# Label-free GAG disaccharide analysis by HILIC-MS/MS for studying diverse biological sample types

**DOI:** 10.64898/2026.04.28.721356

**Authors:** H Davies-Strickleton, George Taylor, James Allsey, Samie Dalgarno, Megan J. Priestley, Isobelle Blair, Nabina Pun, Emily Williams, Magnus Nørregaard Nissen Grønset, Rebecca L. Miller, David Knight, Douglas P. Dyer

**Affiliations:** Discovery Research Platform, Manchester Cell-Matrix Centre, Faculty of Biology, Medicine and Health, Manchester Academic Health Science Centre, University of Manchester; Manchester, UK; Biological Mass Spectrometry Core Facility, Faculty of Biology, Medicine and Health, Manchester Academic Health Science Centre, University of Manchester, Manchester, UK; Manchester Cell-Matrix Centre, Lydia Becker Institute of Immunology and Inflammation, Faculty of Biology, Medicine and Health, Manchester Academic Health Science Centre, University of Manchester; Manchester, UK; Koch Institute for Integrative Cancer Research, Massachusetts Institute of Technology; Boston, MA, USA; Centre for Immuno-Oncology, Nuffield Department of Medicine, University of Oxford, Oxford, UK; Manchester Cell-Matrix Centre, Division of Cell-Matrix Biology and Regenerative Medicine, School of Biological Sciences, Faculty of Biology Medicine and Health, Manchester Academic Health Science Centre, The University of Manchester, Manchester, UK; Copenhagen Center for Glycocalyx Research, Department of Cellular and Molecular Medicine, Faculty of Health Sciences, University of Copenhagen, Blegdamsvej 3, DK-2200 Copenhagen N, Denmark

## Abstract

The extracellular matrix (ECM) and cell surface glycocalyx are key components of biology and play crucial roles in development and tissue function, as well as disease. Proteoglycans, and their glycosaminoglycan (GAG) side chains, are critical components of the ECM and the glycocalyx. GAGs can bind to many different proteins, such as chemokines, and form hydrated barriers around cells. Existing and new methods are helping us to uncover more about the roles of GAGs in biology. Here, we expand on existing technologies and provide streamlined, standardised and well-documented methods that can be easily adopted in standard analytical facilities. We provide extensive detailed step-by-step guides describing sample disruption, GAG disaccharide preparation from biological tissues and their analysis by HILIC-MS/MS. In addition, we demonstrate utility of this method when using a range of different samples as biological sources. This method will sit alongside existing and new techniques to help improve access to GAG analysis, and thereby further the field of understanding GAG function in complex biological contexts.

## Introduction

The extracellular matrix (ECM) and cell surface glycocalyx (the pericellular matrix that surrounds cells) are important structures across biological contexts from development of tissues to removal of invasive pathogens^1–4^. Proteoglycans, which are present on all mammalian cells to different extents, are one of the key components of the ECM and glycocalyx. Proteoglycans have crucial roles in tissue development and function^5^, but also in inflammatory based diseases^2^ and cancer^6^. Classically their function has been primarily focused on their role in forming the basement membrane, underlying endothelial and epithelial cell structures, and as part of the interstitial matrix, that is key to tissue formation. More recent studies have also localised key proteoglycan function to their presence on the cell surface, e.g. endothelial^7^ and immune cells^8^, as part of the cell surface glycocalyx^1^.

Proteoglycans are formed by protein cores that can be secreted or embedded into the cell membrane (e.g. syndecans) that have glycosaminoglycan (GAG) side chains attached^9^. There are a number of different GAGs based on their sequence including, heparan sulfate (HS), chondroitin sulfate (CS) and dermatan sulfate (DS), in addition to the non-sulfated GAG hyaluronan (HA) (Figure 1A)^4^. HS is composed of repeating units of glucosamine (GlcN) and glucuronic/iduronic acid (GlcA/IdoA). CS chains are composed of repeating units of N-acetyl galactosamine (GalNAc) and GlcA, whilst DS incorporates GalNAc, IdoA and GlcA. HS, CS and DS structures are also differentially sulfated during their synthesis to produce charge, and potentially high degrees of chain specificity (Figure 1A). HA, on the other hand, is not protein anchored and is composed of repeating units of N-acetyl glucosamine (GlcNAc) and GlcA.

**Figure 1.**
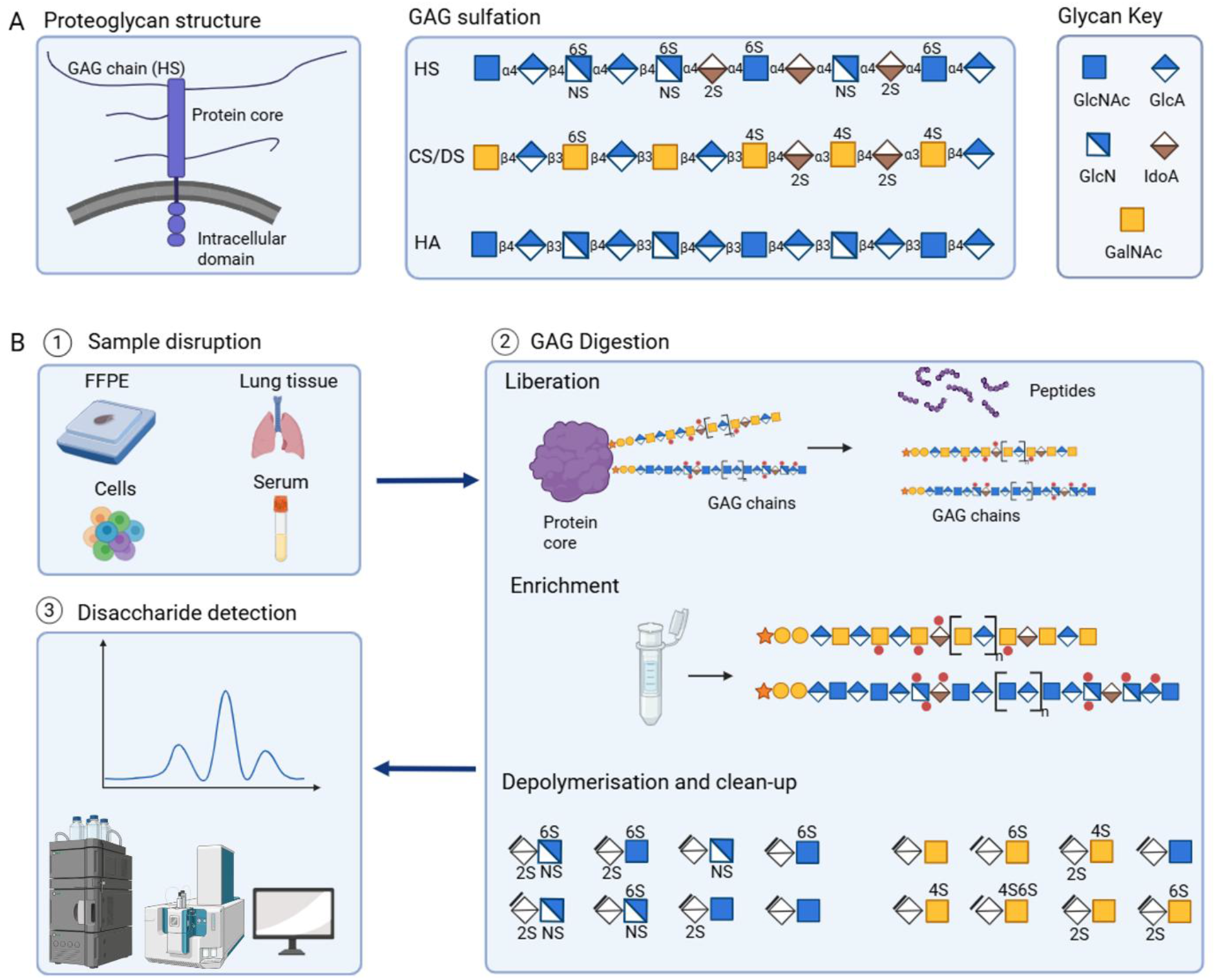
Overview of label-free GAG disaccharide analysis. (A) HS is composed of repeating units of glucosamine (GlcN) and glucuronic/iduronic acid (GlcA/IdoA). CS chains are composed of repeating units of N-acetyl galactosamine (GalNAc) and GlcA, whilst DS incorporates GalNAc, IdoA and GlcA. HA is composed of repeating units of N-acetyl glucosamine (GlcNAc) and GlcA. HS, CS and DS are highly sulfated, whilst HA is non-sulfated. (B) Schematic overview of label-free GAG disaccharide analysis. Sample disruption of diverse biological samples is followed by GAG liberation, enrichment and depolymerisation to yield disaccharides. Label-free disaccharides are analysed by HILIC-MS/MS. Figure created with BioRender.com.

The sulfated GAGs are critical to proteoglycan function due to their large size, hydrophillic nature and negative charge. These properties of the GAG produce hydrated adhesion-resistant structures that form barriers on the cell surface^10^ and regulate their interaction with an array of proteins, including growth factors and chemotactic cytokines^11,12^. Despite their crucial biological function, the study of GAGs in complex biology has been limited by a lack of accessible tools for analysis in biologically relevant contexts.

Recently a number of new approaches have been developed to counter this problem. This includes the establishment of bump-hole engineering to allow “click-chemistry” based labelling of proteoglycans *in vitro*^13^. Furthermore, ToF-SIMS mass spectrometry is also in development to provide spatial information for GAG analysis using specialised apparatus^14^. Alongside these new developments, the classic approach to analyse GAGs from biological samples has been to purify them followed by treatment with GAG lyases to digest their chains into disaccharides, which can then be analysed with or without labelling (e.g. AMAC, 2-AB, Bodipy, Aniline) using several analytical techniques, such as high-performance liquid chromatography (HPLC) or LC-MS^15–27^. This approach has also been evolved to utilise hydrophilic interaction liquid chromatography (HILIC) to increase separation^25,28^. Such tools provide crucial insight into GAG structure, yet they often require chemical labelling with compounds not widely accessible to research labs and/or with limited use due to safety concerns, particularly within academic analytical facilities in the UK.

As part of the Discovery Research Platform for Cell-Matrix Biology, our goal is to facilitate the adoption of diverse analytical methodologies with application to matrix research. By streamlining, standardising and documenting applicable methodologies, methods should be robust, repeatable and easily adoptable by standard analytical facilities. Here, we build on existing published methods^15–28^ and present a comprehensive set of methods for the generation of label-free disaccharides of HS and CS/DS/HA GAGs from tissues, cells and plasma samples, and their analysis by HILIC-MS/MS. We provide links to a collection of detailed step-by-step protocols (https://dx.doi.org/10.17504/protocols.io.x54v956n1l3e/v1), covering everything from disruption of biological material to GAG disaccharide preparation to data analysis. We applied these methods to the analysis of limited amounts of starting biological material from a range of different biological contexts.

## Materials and Methods

### Methods

Detailed step-by-step protocols of the methods in this publication are available as a collection at https://dx.doi.org/10.17504/protocols.io.x54v956n1l3e/v1.

### Materials

GAG disaccharide standards and full-length GAGs were purchased from Iduron, Manchester, UK. Heparinases I, II and III were purchased from New England Biolabs. All other reagents were purchased from Sigma-Aldrich, except the following: calcium chloride (EMD Millipore), ammonium acetate (Avantor), magnesium chloride (Fisher Ccientific), acetonitrile (Fisher Scientific).

### Biological material and disruption

A guide for sample disruption and amount of starting material is available at https://dx.doi.org/10.17504/protocols.io.5jyl88pwrl2w/v1.

Mouse lung tissue was harvested, stored briefly at −20°C and then processed with a Covaris CryoPREP dry pulveriser. The powdered tissue was reconstituted at 200 mg/mL in PBS and homogenised further on a Qiagen Tissue Lyser II with 5 mm stainless steel bead for 4 × 30 seconds, and supernatant used as mouse lung tissue homogenate for GAG disaccharide analysis.

CHO-ZN GS-/-WT cells were maintained in suspension culture under constant agitation (170 rpm) at 37 °C and 5% CO_2_. The cells were in serum-free culture media using a 1:1 mix of BalanCD CHO Growth A (Irvine scientific) and EX-CELL® CD CHO Fusion (Sigma-Aldrich) media, with 2 mM GlutaMAX (GIBCO). Engineered CHO cells were cultured to a concentration of approximately 2 × 10^6^ cells/mL, before cells were centrifugated at 500 x g for 3 mins, washed and centrifuged two times with PBS and stored as cell pellets at −80 °C. After thawing, cell pellets were reconstituted in 50 mM ammonium acetate pH 7.5 in a sonication bath for 20 minutes before preparation of GAG disaccharides.

Mouse kidneys were harvested from an individual C57Bl/6NTac WT male aged 19 weeks, one was snap frozen in liquid nitrogen and stored at −80 °C and the other was fixed with formalin and paraffin embedded (FFPE). The frozen kidney was processed on a Covaris CryoPREP dry pulveriser and ~ 5 mg of powered tissue used per sample replicate. FFPE tissue was sectioned into scrolls (25 µm thick curls), placed inside a 2 mL microfuge tube with a 5 mm stainless steel ball bearing and homogenised on a Qiagen Tissue Lyser II with 200 µL PBS for 4 × 30 seconds. Homogenised FFPE material was decrosslinked at 98 °C for 30 minutes. The homogenisation and decrosslinking steps were repeated once and samples spun at 300 x g for 1 minute. The homogenised tissue was recovered in the PBS by pipetting and was used for GAG disaccharide analysis.

The mouse psoriasis model is described elsewhere^8^. Blood samples were taken from mice by cardiac puncture under terminal anaesthesia and left on ice to coagulate for 4 hours. Blood was centrifuged at 15,000 x g for 10 mins at 4 °C and supernatants collected and frozen prior to HS analysis.

For colon irradiation studies, female C57BL/6 mice (Charles River), aged 11 weeks were irradiated as established previously^29^. Mice were sacrificed 7 days after treatment and the outer colonic mucus layer was collected via a 2 mL PBS wash, and rocked at 4 °C for 5 hours before storage at −80 °C. Samples were thawed, centrifuged and 1 mL supernatant taken for CS/DS/HA analysis.

### Label-free GAG disaccharide generation from biological samples

Detailed step-by-step protocols for label-free GAG disaccharide generation, and vialling for HILIC-MS/MS analysis, are available at https://dx.doi.org/10.17504/protocols.io.kqdg31yxel25/v1 and https://dx.doi.org/10.17504/protocols.io.3byl46progo5/v1, respectively.

Briefly, biological material was digested with 1 mg/mL pronase (with 50 mM ammonium acetate and 1 µM calcium chloride) overnight at 37 °C, and heat inactivated at 98 °C for 10 minutes. DNase I was added at a final concentration of 0.4 mg/mL (with 0.5 µM magnesium chloride) and incubated at 37 °C for 1 hour. Samples were centrifuged and the supernatant filtered through Amicon 500 µL 3 kDa MWCO filters. The retentate (containing full-length GAGs liberated from proteins and DNA) was washed with ddH_2_O and concentrated to ~ 50 µL. For HS disaccharide generation, samples were digested with 1 µL of Heparinase I (12U), II (4U), III (0.7U) enzymes (with ~ 40 mM ammonium acetate pH 7, 3 mM calcium chloride) and incubated at 30 °C overnight (units are given as per the NEB guidelines). For CS/DS/HA disaccharide generation, samples were digested with 10 mU Chondrotinase ABC and incubated overnight at 37 °C. Samples were centrifuged and spiked with 8 ng of an internal disaccharide standard (ΔUA-2S GlcNCOEt-6S), in order to correct for variability in the extraction and analysis from this point. Samples were passed over an OASIS HLB SPE to remove hydrophobic contaminants. Disaccharides were lyophilised, reconstituted in 20 µL water and diluted 1 in 20 into 75% acetonitrile, 44 mM ammonium acetate for HILIC-MS/MS analysis (this provided an internal standard concentration of 20 ng/mL). For each batch of sample preparation, a positive control sample QC (e.g. lung tissue homogenate) was put through the entire protocol to ensure expected data were obtained.

### Equimolar disaccharide standard preparation

For each type of GAG (HS or CS/DS/HA) an equimolar disaccharide standard (a mixture of disaccharide standards at equal molar concentration, as defined by their UV absorption at 232 nm) was prepared as described further: https://dx.doi.org/10.17504/protocols.io.n92ld6knxg5b/v1. The purpose of this standard was to correct for disaccharide specific variability in analysis and to provide a single point calibration for the approximation of disaccharide amount (semi-quantification). Absolute quantification would require multi-point calibration and validation within matrices, however, the semi-quantification presented here is appropriate for the relative quantification of disaccharide amounts, total GAG amounts (by summation of disaccharide amounts) and disaccharide composition (relative ratio of disaccharides compared to total GAG).

In brief, each individual disaccharide standard was reconstituted using ddH_2_O and the UV absorbance at 232 nm was determined. Correction factors were calculated by dividing the UV absorbance of each standard by the UV absorbance obtained from HD002 (HS) or CD001 (CS/DS/HA). HD002 and CD001 were selected as the reference standards due to their consistency in an initial multi-batch comparison. Corrected standard concentrations were used to generate a single equimolar stock solution consisting of 100 µM of each disaccharide standard, which was subsequently diluted to 1 µM in ddH_2_O.

The 1 µM equimolar stock solution was diluted 1 in 20 in 75 % acetonitrile, 44 mM ammonium acetate and spiked with internal standard at a final concentration of 20 ng/mL (ΔUA-2S GlcNCOEt-6S - to correct for variability in instrument performance)). This provided a 50 nM equimolar disaccharide standard, that when back-calculated provided the equivalent of 20 pmol of each disaccharide in the extracted sample, and was then used to calculate an in-sample amount.

### HILIC-MS/MS data acquisition and data processing of GAG disaccharides

A detailed step-by-step protocol for data acquisition and processing is available at https://dx.doi.org/10.17504/protocols.io.n92ld6keog5b/v1.

HILIC-MS/MS was carried out on a Shimadzu Nexera LC 30 AD coupled to a SCIEX 7600 ZenoTOF mass spectrometer. Separations were performed on 5 µL injections at 0.25 mL/min using an InfinityLab Poroshell 120 HILIC-Z column with dimensions of 150 × 2.1 mm, 2.7 μm particle size equipped with a guard column, and mobile phase A (25 mM ammonium acetate in water) and B (25 mM ammonium acetate in 85% acetonitrile). A column temperature of 50 °C and autosampler temperature of 5 °C were used. HS disaccharides were eluted on an LC gradient of 94-90 % B in 25 min, which was achieved by running solvents at 94-94-90-5-5-94-94 % B in 0-3-25-25.1-27-27.1-30 minutes. CS/DS/HA disaccharides were eluted on an LC gradient of 98-87 % B in 25 minutes, which was achieved by running solvents at 98-98-87-87-5-5-98-98 % B in 0-16-24-26-26.2-30-30.1-35 minutes. New columns were observed to require LC gradients to be adjusted by +/− 1-5 % B.

MS data were acquired in negative mode (curtain gas, 30 psi; ion spray voltage, −4500 V; temperature, 250 °C; ESI nebuliser gas pressure, 50 psi; heater gas pressure, 70 psi). High resolution-multiple reaction monitoring of the precursors (as shown in Supplementary Tables 1&2) was performed to generate full scan MS/MS data. For data processing, fragment ions to be used for quantification were selected based on their specificity at the selected retention time and their positive annotation in GlycoWorkbench^30^ (fragment ions are listed in Supplementary Tables 1 & 2). Quantification was performed in SCIEX OS (V3.0), and summed peak area ratios (relative to the spiked internal disaccharide standard) were calculated. Each disaccharide peak area ratio was normalised to that of the equimolar disaccharide standard, to correct for disaccharide specific variability in analysis and provide a single point calibration for the approximation of disaccharide amount in the extracted sample (semi-quantification). The value is considered as semi-quantitative as (a) comparison is to a single calibration point and (b) full validation within each biological matrix has not been performed. The semi-quantification presented here is appropriate for the relative quantification of disaccharide amounts, total GAG amounts (by summation of disaccharide amounts) and disaccharide composition (relative ratio of disaccharides compared to total GAG).

## Results

### Label-free, non-toxic, GAG disaccharide generation

Given the recent breakthroughs in understanding the wider biological functions of GAGs, we endeavoured to develop a streamlined and standardised method, supported by extensive step-by-step guides (https://dx.doi.org/10.17504/protocols.io.x54v956n1l3e/v1) and applicability to a wide range of biologically relevant samples, such as tissue biopsies, FFPE tissue, cells, and plasma. The methodology employs sample disruption, followed by GAG liberation, enrichment and depolymerisation into label-free disaccharides, which are then analysed by HILIC-MS/MS (Figure 1).

The initial sample disruption was tailored to the sample type and utilised tools including bead homogenisation of FFPE tissue, dry pulverisation of frozen tissue via cryoPREP and sonication for cell lysis. Following disruption, lysed samples were digested with pronase for the enzymatic degradation of proteins into peptides, and DNase for the digestion of DNA into nucleotides. The resulting small peptides and nucleotides, could then be separated from the large GAGs by size separation with a molecular weight cut off (MWCO) filter and centrifugation. Enriched GAGs were depolymerised with heparinases and/or chondroitinase ABC digestion, as per traditional ways of generating HS and CS/DS/HA disaccharides, respectively^31^. Here, GAG disaccharides remained label-free, to reduce further sample processing and labelling inefficiencies, and were cleaned-up by solid phase extraction prior to HILIC-MS/MS analysis.

### HILIC-MS/MS detection of label-free GAG disaccharides

For the detection of label-free GAG disaccharides, HILIC was employed based on its ability to separate highly polar compounds and was coupled to tandem mass spectrometry to provide specificity and sensitivity. Pure GAG disaccharide standards (Iduron, UK) were used to develop HILIC-MS/MS conditions for the unique detection of each disaccharide. HILIC chromatographically resolved many of the disaccharides, including the HA disaccharide ΔUA-(ß1-3)-GlcNAc and its structural isomer, the unsulfated CS/DS disaccharide ΔUA(1-4)-GalNAc (Figure 2). Whilst isomeric disaccharides that did not resolve chromatographically were detected by unique MS/MS fragments (Supplementary Tables 1&2). Where multiple unique fragments were available, they were summed in order to increase sensitivity (Supplementary Tables 1&2). HS and CS/DS/HA were run as separate data acquisition methods based on overlapping retention times and lack of unique MS/MS fragments.

**Figure 2.**
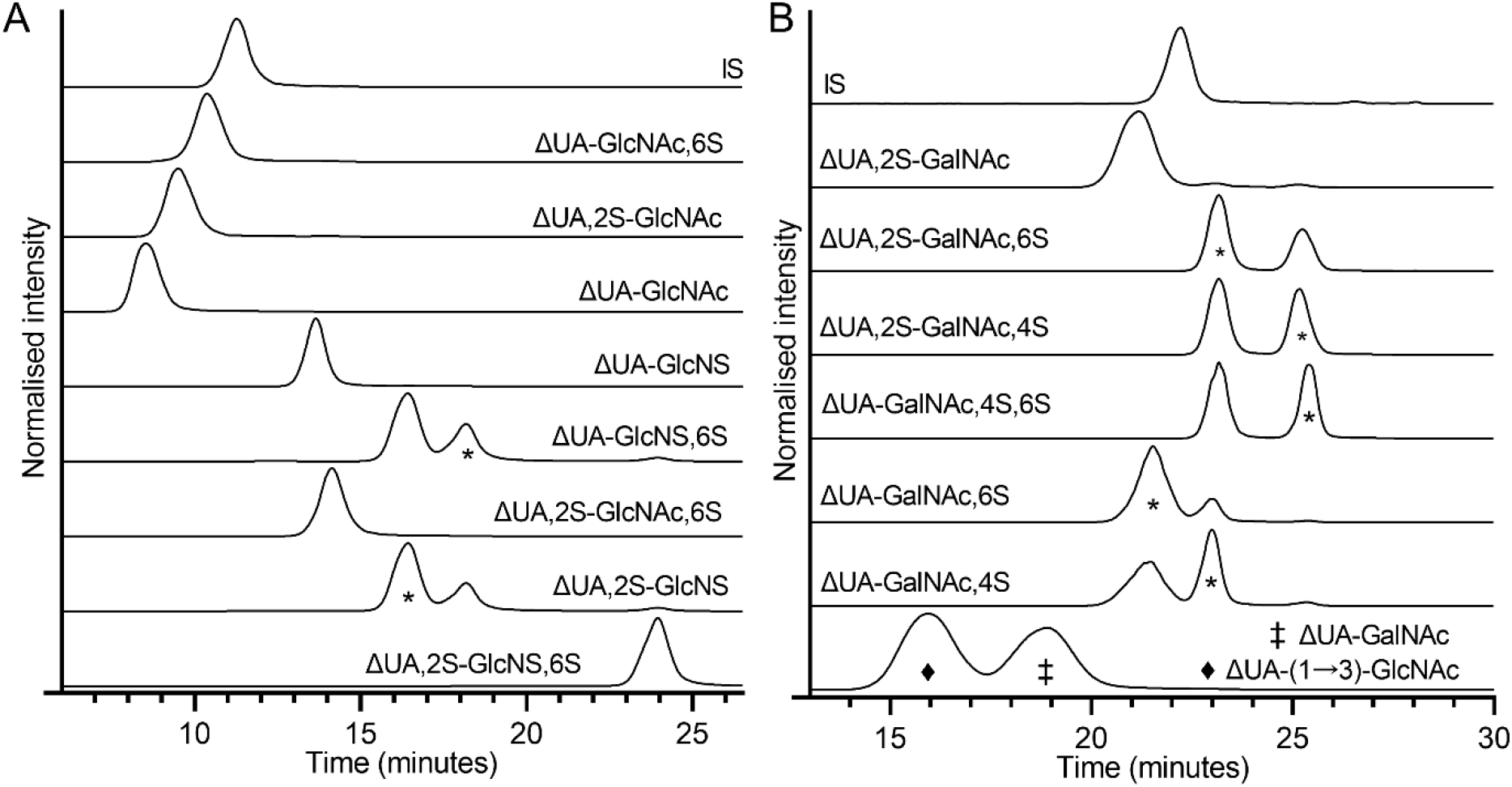
HILIC-MS/MS of GAG disaccharide standards. Representative extracted ion chromatograms of (A) HS and (B) CS/DS/HA disaccharides by HILIC-MS/MS analysis. Symbols denote the analyte peak, additional peaks were due to isomers.

Peak area ratios (relative to the internal standard peak area) were then calculated for each disaccharide. The internal standard (the synthetic disaccharide ΔUA-2S GlcNCOEt-6S, Iduron, UK, also used elsewhere^32,33^), served to account variability in sample processing and instrument performance. The peak area ratios were normalised to that of an equimolar disaccharide standard (an equimolar mixture of HS or CS/DS/HA disaccharides). This step corrected for disaccharide specific variability in analysis (e.g. ionisation efficiencies) and provided a single point calibration to approximate disaccharide amounts (semi-quantification). This approach is appropriate for the relative quantification of disaccharide amounts and total GAG amounts (by summation of disaccharide amounts), and the calculation of disaccharide composition (relative ratio of disaccharides compared to total GAG). Thus, providing an efficient tool for the exploration of GAG changes in a variety of biological contexts.

### Robust GAG disaccharide analysis of low amounts of diverse sample types

Given the need to determine the function of GAGs across different biological contexts (e.g. tissues and diseases) we used the label-free GAG disaccharide preparation and HILIC-MS/MS analysis to investigate GAG profiling of diverse sample types. Robustness was first assessed by measurement of samples of mouse lung tissue homogenate prepared in multiple replicates (Figure 3, Supplementary Table 3; intra-sample precision). Disaccharide amounts and total GAG amounts demonstrated low Coefficient of Variation (CV) of 2-30 %, indicating high consistency and intra-sample precision. For disaccharide composition (relative ratio of disaccharides compared to total GAG), intra-sample precision was also high, with CV of 1-18 %. Therefore, the method has the necessary robustness for the routine analysis of biological samples.

**Figure 3:**
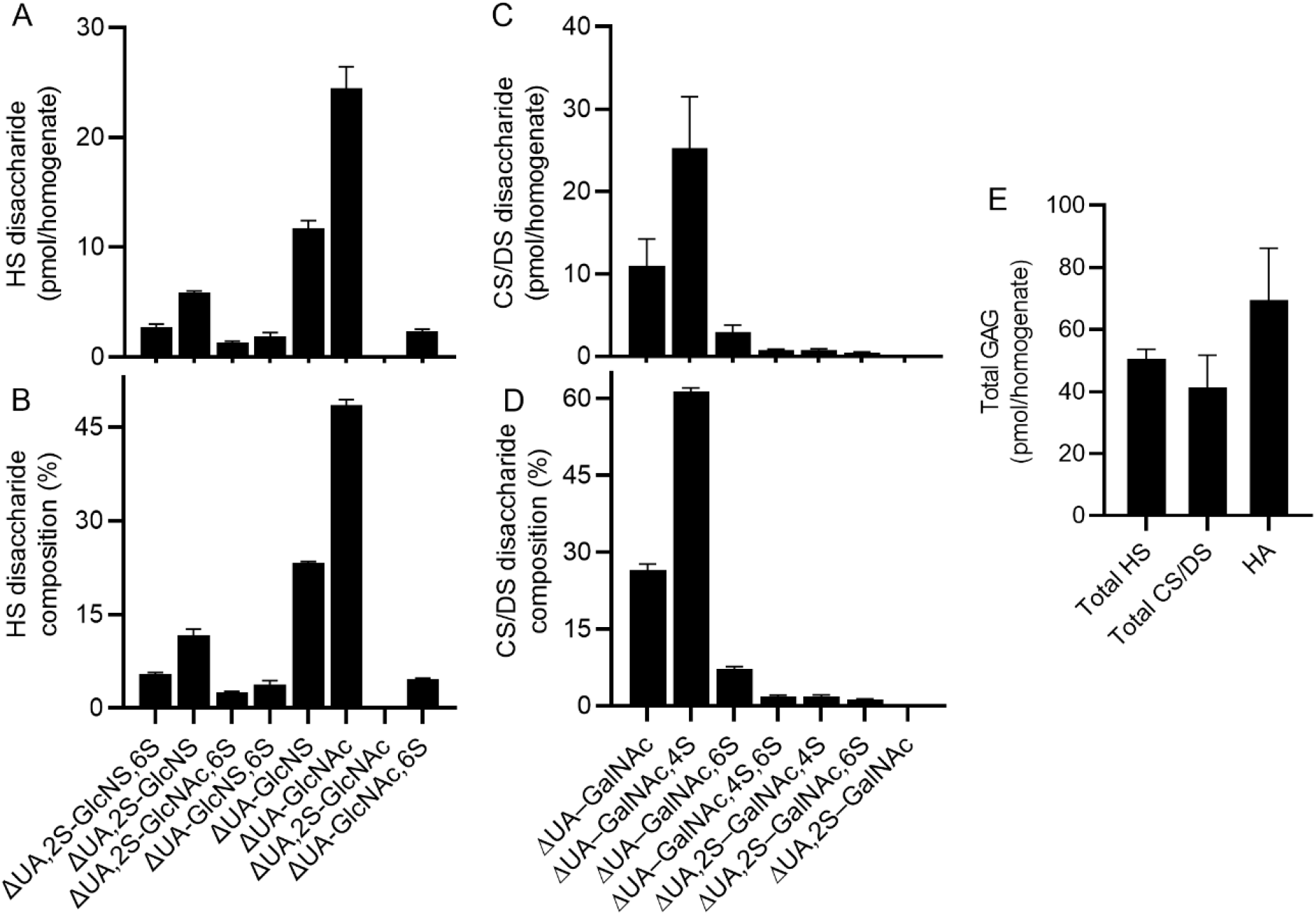
Robust detection of GAG disaccharides from tissue homogenates. Samples of mouse lung tissue homogenate were prepared in replicate and analysed by HILIC-MS/MS for HS disaccharides (A, B) and CS/DS/HA disaccharides (C, D). Disaccharide amount and total GAG amount were determined (A, C, E), and disaccharide compositions calculated (B, D). (n=5 replicates of the same sample, measured in a single experiment; intra-sample precision).

Next, CHO cell GAG chains, which have been well characterised previously^31,34,35^, were analysed to assess the consistency of disaccharide detection across varying amounts of starting material. The disaccharide composition was in good agreement with previous reports using traditional HPLC-FLD methodologies for both HS and CS/DS disaccharides^31^ (Supplementary Table 4, data from 1e^6^ cells). As the number of cells per sample was reduced, the HS disaccharide composition remained largely consistent down to 2e^4^ cells (Figure 4A). In contrast, the CS/DS disaccharide composition was dominated by a single disaccharide (93 % ΔUA-GalNAc,4S, based on data from 1e^6^ cells) and the less abundant disaccharides were not detected in samples of cell number below 1e^6^ cells, yielding disaccharide compositions of 100 % ΔUA-GalNAc,4S for samples of lower cell numbers (2e^4^ – 1e^5^ cells, Figure 4B). Despite this, across samples of 2e^4^ – 1e^6^ cells, the disaccharide amounts and total GAG amounts scaled linearly with the number of cells per sample for both HS (r^2^ > 0.99) and CS/DS (r^2^ > 0.99) data (Figure 4C-F). HA was not detected. Together, the CHO cell data demonstrate that the method can reproduce information from GAG sulfation for biological samples reported elsewhere, can provide relative comparison of total GAG amounts (and the most abundant individual disaccharides) from different amounts of starting material, and can utilise even low amounts of starting material.

**Figure 4:**
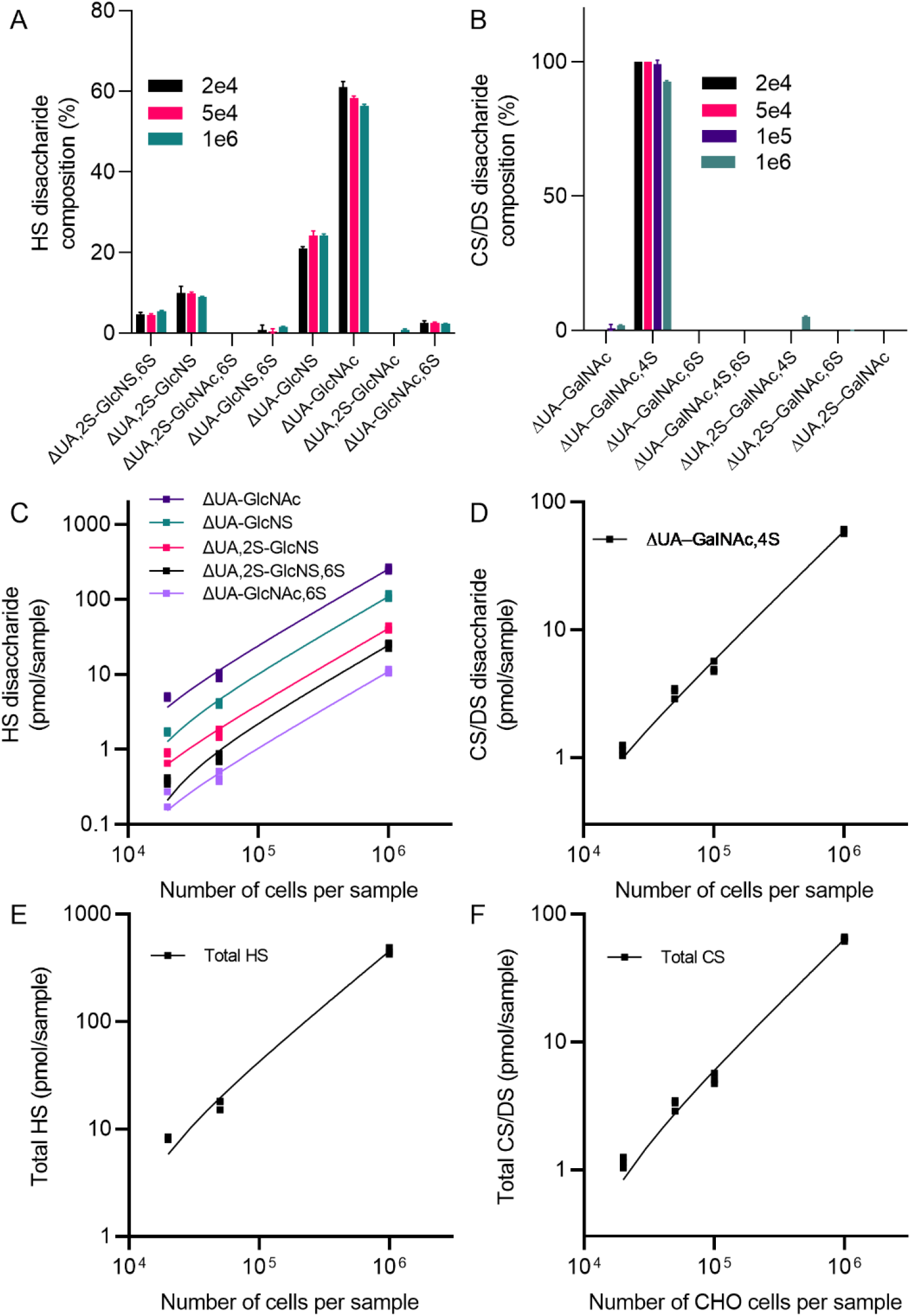
GAG analysis of WT CHO cells correlates with the number of cells analysed. Disaccharide composition for HS (A) and CS/DS (B) from different starting numbers of CHO WT cells: 2e^4^ (black), 5e^4^ (pink), 1e^5^ (blue, CS/DS only) and 1e^6^ (green) CHO cells (n=3 replicates per sample group). (C) HS and (D) CS/DS/HA disaccharide amount detected (pmol/sample) from different starting numbers of CHO cells. Data are shown for disaccharides detected in all sample groups; for CS/DS disaccharides this was only ΔUA-GalNAc. (E) HS and (F) CS total GAG detected (pmol/sample) from different starting numbers of CHO cells. All data (C-F) were fitted with a simple linear regression and gave r^2^ > 0.99.

To assess application of the method to preserved biological samples, mouse kidneys were harvested from a single animal, with one frozen and one preserved by FFPE. Frozen tissue was excised and FFPE scrolls were homogenised, and GAG disaccharides prepared and analysed. Disaccharides from both HS and CS/DS were detected in both sample types (Figure 5, Supplementary Table 5). The amount of HS and CS/DS was lower in FFPE tissue compared to frozen tissue, whilst HA was only detectable in frozen tissue. This most likely reflects the lower amount of starting material in the FFPE scroll samples (3 × 25 µm sections) compared to frozen tissue (5 mg tissue biopsy) analysed here. Disaccharide composition for HS and CS/DS were highly consistent across FFPE and frozen tissue and in agreement with previous analysis^36^. Thus, these data demonstrate that the method can assess samples of different preservation type.

**Figure 5:**
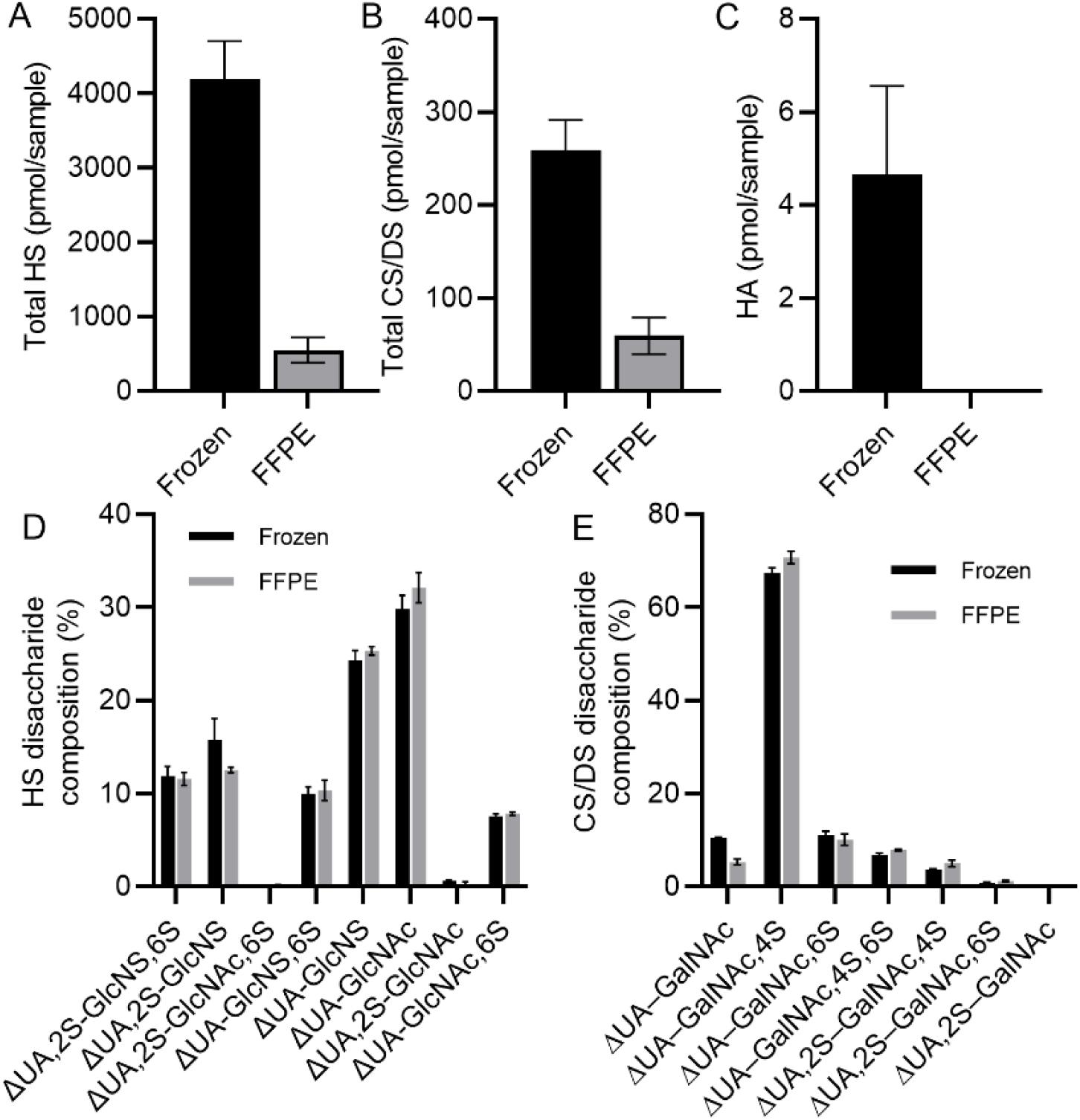
Analysis of differentially preserved biological samples. GAG disaccharide analysis was performed on frozen (black) and FFPE (grey) mouse kidney tissue from an individual animal. Total HS (A), total CS/DS (B) and total HA (C) amounts were determined, as well as disaccharide compositions of HS (D) and CS/DS (E). N=5 technical replicates.

### Differences in GAGs revealed across biological samples

Next, the ability of the method to reveal new biological insight was explored. We have previously demonstrated that HS levels are elevated in the serum of mice in a model of psoriasis using ELISA approaches^8^. We analysed the same samples with our method and confirmed increased circulating HS in the inflamed mouse serum (Figure 6A). Furthermore, the analysis here suggests that this change was largely due to an increase in the unsulfated HS disaccharide ΔUA-GlcNAc. CS/DS disaccharide analysis of the same samples also indicated an increase in the unsulfated CS/DS disaccharide ΔUA-GalNAc, and a slight increase in total CS/DS amounts, although the latter was not statistically significant (Figure 6B). The amount of HA did not increase (Figure 6B).

**Figure 6:**
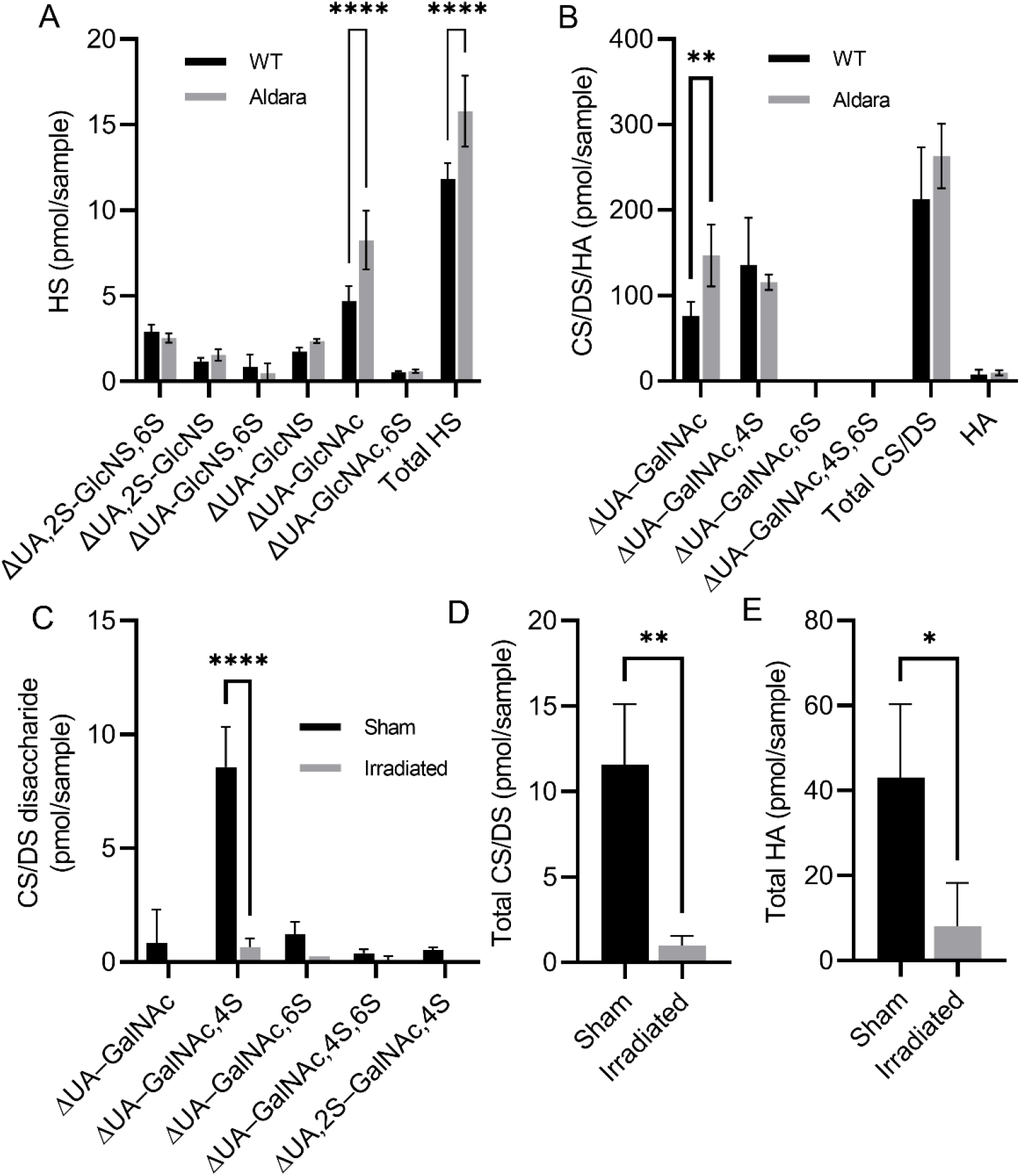
GAG changes in biological models of disease. In a mouse model of psoriasis, (A) HS and (B) CS/DS/HA disaccharides differences were detected between untreated (black) and treated (grey) mice (n=4 biological replicates, two-way anova). (C&D) GAG changes in colonic mouse mucus after sham treatment (black) or radiation (grey) for (C) CS/DS disaccharides and (D) total CS/DS and HA amounts (n=3 biological replicates, unpaired T-test).

In another application, GAG biology was shown to change in mouse colonic mucus after irradiation to model normal tissue toxicity induced by abdominal radiotherapy (Figure 6&D). This revealed a drastic reduction in the amounts of HA and CS compared to sham treatment.

## Discussion

Here, we present a streamlined and standardised detailed step-by-step method for the preparation of label-free GAG disaccharides from low amounts of diverse biological samples, their analysis by HILIC-TOF/MSMS and application to identify changes to GAGs in biologically relevant contexts.

We provide extensive step-by-step guides for each stage, enabling reproduction of the methods in their entirety. To further support method reproducibility and accessibility, we utilised sample preparation strategies with commercially available consumables, such as MWCO filters. Additionally, disaccharides were kept label-free, which served to both reduce preparation time and bypass health and safety barriers to utilising hazardous chemicals involved in some labelling strategies. Thus, further enhancing accessibility of this method, particularly in UK institutions and universities where health and safety barriers limit alternative approaches.

The HILIC-MS/MS employed here for detection of the label-free disaccharides provided the means to detect each GAG disaccharide via unique MS/MS fragmentation and/or chromatographic separation. The fragments identified here agree with those identified previously^37,38^. The overlapping HILIC retention of HS and CS/DS/HA disaccharides and lack of unique MS/MS fragments meant that the method required separate heparinase/chondroitinase ABC digestion. Whilst this is a common approach for GAG disaccharide analysis elsewhere^15^, other groups have shown that AMAC^39,40^ or 2AB-labelled GAG disaccharide approaches can be run with dual digest and a single LC-MS/MS method^41^.

The approach taken here was to utilise a single calibration point for approximation of disaccharide amounts (semi-quantification). Alternative methods for quantification include the utilisation of 13C-labelled disaccharide and polysaccharides^25,42^. The quantification performed here was appropriate for the relative comparison of disaccharide amounts, total GAG amounts and calculation of disaccharide composition, and was shown to be applicable to diverse sample types and a range of starting material including those in limited quantity (e.g. 2e^4^ cells (Figure 4), 3 x FFPE scrolls and < 5 mg tissue (Figure 5).

We have used this method to analyse low sample sizes, producing a sensitivity that is comparable to other approaches^43^. Method robustness and accuracy were also demonstrated via consistency of disaccharide amounts and disaccharide composition from replicate samples of lung tissue homogenate (Figure 3), and consistency of disaccharide composition of CHO cells with established methods utilising labelled GAG disaccharides and HPLC-fluorescent detection (FLD) analysis elsewhere (Figure 4 and Supplementary Table 3)^31,34,35^, respectively. In addition, GAG levels correlated well with cell numbers (r^2^=0.99; Figure 4), demonstrating the ability of the method to reliably deduce differences in GAG disaccharide and total GAG levels from different amounts of starting material. Thus, this method offers a robust and reliable tool to study different biological contexts. This was further exemplified here by demonstrating conformation of differences in models of inflammatory disease and new observations in a tissue toxicity after irradiation model (Figure 6). By studying diverse sample types, this analysis can be applied to a broad range of research contexts and should help to advance the study of GAGs in complex biological contexts.

We anticipate that this method will combine with existing LC-MS approaches and newly developed methods, such as ToF-SIMS spatial GAG mass spectrometry, to give context to the use of existing, and improving, antibody-based detection of GAGs. Together this combination of tools will help to improve our understanding of GAG function in complex biological contexts.

## Acknowledgements

We thank Becky Dodd and Judi Allen (Manchester Cell-matrix Centre, University of Manchester, UK) for the use of the Qiagen TissueLyser II.

## Funding

Sir Henry Dale fellowship jointly funded by the Wellcome Trust and Royal Society 218570/Z/19/Z (DD)

Wellcome Trust Career Development Award 319823/Z/24/Z (DD)

Wellcome Trust Discovery Research Platform 226804/Z/22/Z (DD and HDS)

The Novo Nordisk Foundation grant NNF22OC0073736 (to RLM); The Human Frontier Science Program grant RGEC31/2023 (to RLM); The Innovations Fonden grant 4295-00042B (to RLM).

## Author Contributions

Conceptualization: HDS, RLM, DK, DPD

Methodology: HDS, GT, JA, SD, RLM, DK, DPD

Investigation: HDS, GT, JA, SD, MJP, IB, NP, MNNG, RLM, DK, DPD

Visualization: HDS, SD, DK, DPD

Resources: HDS, GT, JA, SD, MJP, IB, NP, MNNG

Funding acquisition: RPM, DK, DPD

Project administration: HDS, DK, DPD

Supervision: RLM, DK, DPD

Writing – original draft: HDS, DK, DPD

Writing – review & editing: HDS, GT, SD, MJP, IB, NP, MNNG, RLM, DK, DPD

## Data and materials availability

Any data not explicitly evident in the paper will be available on reasonable request.

