## Supplementary information for "Label-free GAG disaccharide analysis by HILIC-MS/MS for studying diverse biological sample types"

**Affiliations:**

**Supplementary Table 1: List of ion transitions (precursors and fragment ions) for HS disaccharide analysis**

| Analyte Name | Relative Retention Time | Precursor (Q1) Mass (Da) | Fragment (Q3) Mass (Da) |
|---|---|---|---|
| Internal standard | 1 | 552.0336 | 472.0781 |
| ΔUA,2S-GlcNS,6S | 2.13 | 287.4784 | 96.9599 |
| | | | 247.5 |
| | | | 137.9865 |
| | | | 357.0129 |
| | | | 157.0141 |
| | | | 287.4784 |
| ΔUA,2S-GlcNS | 1.46 | 247.5 | 96.96 |
| | | | 247.5 |
| | | | 137.9866 |
| | | | 157.0143 |
| | | | 357.0139 |
| ΔUA,2S-GlcNAc,6S | 1.25 | 268.5049 | 96.9602 |
| | | | 157.0143 |
| | | | 138.9711 |
| | | | 370.046 |
| | | | 268.5049 |
| ΔUA-GlcNS,6S | 1.61 | 247.5 | 300.0404 |
| | | | 247.5 |
| | | | 258.0303 |
| | | | 96.96 |
| | | | 157.0146 |
| | | | 137.9866 |
| | | | 357.0139 |
| ΔUA-GlcNS | 1.21 | 416.0514 | 137.9867 |
| | | | 416.0514 |
| | | | 175.0255 |
| ΔUA-GlcNAc | 0.76 | 378.1048 | 378.1048 |
| | | | 175.0251 |
| | | | 157.015 |
| | | | 277.0567 |
| ΔUA,2S-GlcNAc | 0.85 | 458.0617 | 236.9712 |
| ΔUA-GlcNAc,6S | 0.92 | 458.0617 | 175.0256 |
| | | | 357.0148 |
| | | | 300.0403 |

**Supplementary Table 2: List of ion transitions (precursors and fragment ions) for CS/DS/HA disaccharide analysis**

| Analyte Name | Relative Retention Time | Precursor (Q1) Mass (Da) | Fragment (Q3) Mass (Da) |
|---|---|---|---|
| Internal standard | 1 | 552.0336 | 472.0781 |
| ∆UA-(1–3)-GlcNAc | 0.72 | 378.1041 | 175.0256<br>157.0146<br>378.1045 |
| ∆UA-GalNAc | 0.85 | 378.1041 | 175.0249<br>157.0143<br>378.1045 |
| ∆UA-GalNAc,4S | 1.04 | 458.0614 | 300.0399<br>342.0504<br>175.0247<br>157.0143<br>282.029<br>458.0614 |
| ∆UA-GalNAc,6S | 0.97 | 458.0614 | 282.0288<br>175.0248 |
| ∆UA-GalNAc,4S,6S | 1.15 | 268.5053 | 282.0294<br>175.0249 |
| ∆UA,2S-GalNAc,4S | 1.13 | 268.5053 | 300.0398 |
| ∆UA,2S-GalNAc,6S | 1.04 | 268.5053 | 300.0398<br>282.0294<br>157.0146<br>96.96<br>268.5053 |
| ∆UA,2S-GalNAc | 0.95 | 458.0614 | 236.971 |

**Supplementary Table 3: Intra-sample precision of disaccharide amount, total GAG amount and disaccharide composition from lung tissue homogenate replicates (n=5)**

| | | pmol/homogenate | | Disaccharide composition % | |
|---|---|---|---|---|---|
| | | Mean | % CV | Mean | % CV |
| HS disaccharide | ΔUA,2S-GlcNS,6S | 2.76 | 9 | 5.46 | 5 |
| | ΔUA,2S-GlcNS | 5.90 | 2 | 11.71 | 8 |
| | ΔUA,2S-GlcNAc,6S | 1.32 | 10 | 2.60 | 5 |
| | ΔUA-GlcNS,6S | 1.90 | 19 | 3.75 | 18 |
| | ΔUA-GlcNS | 11.77 | 6 | 23.30 | 1 |
| | ΔUA-GlcNAc | 24.53 | 8 | 48.51 | 2 |
| | ΔUA,2S-GlcNAc | BLD | N/A | BLD | N/A |
| | ΔUA-GlcNAc,6S | 2.36 | 8 | 4.66 | 3 |
| CS/DS disaccharide | ΔUA – GalNAc | 11.00 | 30 | 26.47 | 5 |
| | ΔUA – GalNAc, 4S | 25.29 | 25 | 61.41 | 1 |
| | ΔUA – GalNAc, 6S | 2.99 | 26 | 7.25 | 6 |
| | ΔUA – GalNAc, 4S, 6S | 0.75 | 19 | 1.83 | 14 |
| | ΔUA, 2S – GalNAc, 4S | 0.75 | 22 | 1.85 | 18 |
| | ΔUA, 2S – GalNAc, 6S | 0.48 | 17 | 1.18 | 15 |
| | ΔUA, 2S – GalNAc | BLD | N/A | BLD | N/A |
| | Total HS | 50.53 | 6 | N/A | N/A |
| | Total CS/DS | 41.26 | 25 | N/A | N/A |
| | Total HA | 69.51 | 24 | N/A | N/A |

### Supplementary Table 4: Disaccharide composition of CHO cells analysed here compared to literature

| | | Davies-Strickleton et al. 2026 ($1e^6$ cells)* | Chen et al. 2018** |
|---|---|---|---|
| HS disaccharide | ΔUA,2S-GlcNS,6S | 5.4 | 6.0 |
| | ΔUA,2S-GlcNS | 9.1 | 16.0 |
| | ΔUA,2S-GlcNAc,6S | BLD | 0.0 |
| | ΔUA-GlcNS,6S | 1.6 | 0.9 |
| | ΔUA-GlcNS | 24.2 | 20.0 |
| | ΔUA-GlcNAc | 56.4 | 54.0 |
| | ΔUA,2S-GlcNAc | 0.9 | 1.9 |
| | ΔUA-GlcNAc,6S | 2.4 | 1.3 |
| CS/DS disaccharide | ΔUA – GalNAc | 2.0 | 3.5 |
| | ΔUA – GalNAc, 4S | 92.7 | 94.3 |
| | ΔUA – GalNAc, 6S | 0.0 | 0.0 |
| | ΔUA – GalNAc, 4S, 6S | 0.0 | 0.0 |
| | ΔUA, 2S – GalNAc, 4S | 5.2 | 2.2 |
| | ΔUA, 2S – GalNAc, 6S | 0.1 | 0.0 |
| | ΔUA, 2S – GalNAc | 0.0 | 0.0 |

* Data is an average of 5 technical replicate

** Data converted to % from Supplementary Table 3 within Chen et al 2018

**Supplementary Table 5: Disaccharide composition of frozen and FFPE mouse kidney tissue (average from n=5 technical replicates)**

| | | Mean disaccharide composition % | |
|---|---|---|---|
| | | Frozen | FFPE |
| HS disaccharide | ΔUA,2S-GlcNS,6S | 11.9 | 11.6 |
| | ΔUA,2S-GlcNS | 15.8 | 12.5 |
| | ΔUA,2S-GlcNAc,6S | 0.1 | 0.1 |
| | ΔUA-GlcNS,6S | 10.0 | 10.4 |
| | ΔUA-GlcNS | 24.3 | 25.3 |
| | ΔUA-GlcNAc | 29.8 | 32.1 |
| | ΔUA,2S-GlcNAc | 0.6 | 0.2 |
| | ΔUA-GlcNAc,6S | 7.5 | 7.8 |
| CS/DS disaccharide | ΔUA – GalNAc | 10.3 | 5.3 |
| | ΔUA – GalNAc, 4S | 67.3 | 70.7 |
| | ΔUA – GalNAc, 6S | 11.1 | 10.1 |
| | ΔUA – GalNAc, 4S, 6S | 6.8 | 7.8 |
| | ΔUA, 2S – GalNAc, 4S | 3.6 | 5.0 |
| | ΔUA, 2S – GalNAc, 6S | 0.8 | 1.2 |
| | ΔUA, 2S – GalNAc | 0.0 | 0.0 |
